# Angular leaf spot in *Acmella oleracea* caused by a foliar nematode

**DOI:** 10.1101/2023.10.24.563638

**Authors:** Marcela de Freitas Silva, Felipe Castro Faccioli, Amanda Pereira Honório, Andressa Rodrigues Fonseca, Alessandra de Jesus Boari, Dalila Sêni Buonicontro

## Abstract

Jambu plants (*Acmella oleracea*) exhibiting necrotic angular leaf spots were collected in Belém - Pará, Brazil. After previous analysis, the presence of nematodes from *Aphelenchoides* genus was observed. These nematodes were cultured on *Fusarium* sp. and subsequently morphologically and molecularly characterized for species-level identification. *Aphelenchoides* sp. associated with jambu exhibited morphological and morphometric characteristics very similar to those of species within the *A. besseyi* complex (*A. besseyi*, *A. oryzae* and *A. pseudobesseyi*), but these characteristics were not sufficient to separate them into a unique species. The Bayesian inference analysis, utilizing the expansion segment of the large subunit (D2-D3 LSU) of nuclear ribosomal DNA, yielded results with a high posterior probability, indicating that the *Aphelenchoides* sp. associated with jambu belongs to the *A. pseudobesseyi* species. Under controlled conditions, the reproduction of the nematode in the leaf tissues (FR > 1) was observed, resulting in disease symptoms. The highest reproduction rate of *A. pseudobesseyi* (FR = 2.6) was observed from inoculation with 100 nematodes per leaf. It is concluded that *A. pseudobesseyi* is the etiological agent of jambu angular leaf spot. For future research, like evaluating the resistance of jambu to this nematode, it is recommended to inoculate a maximum of 100 nematodes per leaf.

## Introduction

Jambu (*Acmella oleracea*) is a vegetable from South America, belonging to the Asteraceae family, and it is used in traditional cuisine in the Northern region of Brazil. In addition to its gastronomic value, the plant is also used in traditional medicine (Borges et al. 2013; Maria-Ferreira et al. 2014). Several bioactive compounds isolated from *A. oleracea* have anesthetic, anti-inflammatory, antioxidant, and diuretic activity (Freitas-Blanco et al. 2016; Dallazen et al. 2020; 2022; Rondanelli et al. 2020). Its main bioactive compound, spilanthol, has antifungal, bacteriostatic, and insecticidal effects (Spinozzi et al. 2022). This compound has already been reported to control insects of agricultural importance, such as *Helicoverpa zea*, *Plutella xylostella*, and *Tuta absoluta*, and insect vectors of human diseases, including *Aedes aegypti*, *Anopheles* spp. and *Culex quiquefasciatus* (Spinozzi et al. 2022).

Despite the potential of this plant in the gastronomic and pharmaceutical fields, little is known about the pathogens that affect it and may limit its cultivation. The few records found in the literature report the incidence of fungi *Tecaphora spilanthes* and *Puccinia cnici-oleracei*, which cause, respectively, symptoms such as galls and deformations in the stem of the plant and rust (Coutinho et al. 2006; Carmo et al. 2016), in addition to the nematode *Meloidogyne* sp., which causes the formation of root galls (Singh et al. 2000; Homma et al. 2011).

Jambu plants showing necrotic leaf spots were found in Belém city, Pará state, in Brazil. The symptoms observed were distinct from those disease jambu previously reported, in this case, being a disease unknown. After preliminary analysis, nematodes belonging to the *Aphelenchoides* genus were found associated with the leaf spots. Therefore, the aim of this study was to identify *Aphelenchoides* sp. at the species level employing integrative taxonomy and determining if this nematode is the causal agent of the leaf spot observed in jambu.

## Materials and methods

### Obtaining, extracting and cultivating of nematodes

The nematodes were extracted by technique of Coolen and D’Herde (1972) from symptomatic jambu leaves, which were collected in the vegetable field at Federal Rural University of the Amazônia (UFRA), in Belém-PA, in 2019. In order to provide a pure culture, those nematodes were multiplied in *Fusarium solani* and *F. pseudocircinatum* colonies. These fungal species were used to evaluate which one ensured greater nematode multiplication. Thus, approximately 25 specimens of *Aphelenchoides* sp. were hand-picked, with the aid of a stereoscopic microscope, surface-desinfested in antibiotic solution (200 mg / mL of ampicillin and 300 mg / mL of chloramphenicol) for 20 min, and transferred to Petri-dishes containing colonies of *F. solani* or *F. pseudocircinatum* which were grown on potato dextrose agar (PDA) (Favoreto et al., 2011). Plates with fungal colonies were previously incubated for ten days in B.O.D. at 25 °C in the dark. After receiving the nematodes, the plates were incubated in B.O.D. for another 20 days. The multiplied nematodes were washed off the culture plates with distilled water, followed by quantification under a light microscope. The final population from the culture of *F. pseudocircinatum* was ten times higher compared to those obtained from *F. solani.* Thus, we decided to use *F. pseudocircinatum* to multiply the inoculum used for taxonomic studies and pathogenicity tests described below.

### Morphological and morphometric characterization of *Aphelenchoides* sp

Nematodes extracted directly from symptomatic leaves and those multiplied in vitro were heat-killed (55 °C, 5 min), fixed in TAF solution (1% triethanolamine and formalin) in a 1 : 1 ratio (v : v) (Courtney et al. 1955), and stored for 48 h in hermetically sealed glass containers. Next, the specimens were infiltrated using the glycerol-ethanol technique (Seinhorst 1959), followed by mounting permanent slides in glycerin (Oliveira and Wilcken 2016). Nematodes were examined and photographed using an Olympus BX53 light microscope fitted with a digital camera (QImaging MicroPublisher RTV-3.3 Color Camera). The images captured were used for measurements and character morphological documentation, made with aid of the Olympus cellSens Dimension v.1.17 software. Both morphological and morphometric characters were used to compare descriptions of morphologically similar species already reported in the literature.

### Molecular characterization of *Aphelenchoides* sp

The DNA extraction was performed using 10 specimens, by technique of Buonicontro et al. (2018). The DNA fragments of the expansion segment of the large subunit (D2-D3 LSU) were amplified using the primers D2A (5’-ACA AGT ACC GTG AGG GAA AGT TG-3’) and D3B (5’-TCG GAA GGA ACC AGC TAC TA-3’) (Ye et al., 2007). The PCR reactions were prepared in 25 μL final reaction volume containing 12,5 μL of GoTaq^®^ G2 Hot Start Colorless Master Mix (Promega, USA), 0,25 μL (10 μM) of each primer, 1 μL of DNA and 11 μL of nuclease-free water to complete the volume of each reaction. Reactions without DNA were included as a negative control. PCR conditions were as follows: initial denaturation at 95°C for 5 min followed by 35 cycles of denaturation at 94°C for 30 s, annealing at 55°C for 45 s, extension at 72°C for 2 min, with a final extension at 72°C for 10 min. Amplified PCR product was resolved by electrophoresis at 100 v for 60 min in 1% agarose gel, purified using Wizard^®^ SV Gel and PCR Clean-Up System (Promega, USA), according to the protocol provided by the manufacturer. The PCR product was cloned using the pGEM®-T Easy vector (Promega, USA) before the sequencing. Plasmid was transformed into *Escherichia coli* electrocompetent cells (Invitrogen, USA) following the manufacturer’s provided protocol. The blue/white selection was used to select the positive clones. After that, plasmid was extracted using the plasmid miniprep kit GeneJET (ThermoScientific, USA) and sequenced. Sequence obtained in this study was deposited in the GenBank database under accession number: OR541661.

### Sequence alignment and phylogenetic analysis

The newly obtained nucleotide sequence was edited with the DNA Dragon software (Hepperle, 2011). Sequences of other Aphelenchoidea available from public databases were included in the analysis adopted the following selection criteria: sequences published in manuscripts about nematodes taxonomy, with information about host and/or original location. Sequence data were aligned using the M-coffee web server, combining different alignment methods (Moretti et al. 2007). The alignment was edited with Gblocks to eliminate poor alignment and divergent positions, maintaining 68% of original positions (Dereeper et al, 2008). *Aphelenchoides fragariae* was chosen as outgroup taxa based on previously phylogenetic analyzes. The best fitting model of nucleotide evolution was determined using the corrected Akaike information criterion (AICc) with JModeltest version 2.2 (Darriba et al. 2012). The best-fit evolution model HKY+I+G with invgamma-distributed rate was selected and Bayesian inference (BI) analyzes employing a Markov Chain Monte Carlo (MCMC) method were performed using MrBayes on XSEDE 3.2.7a via the CIPRES web portal (https://www.phylo.org/) (Miller et al. 2010). BI analysis was initiated with a random starting tree and was run with four MCMC chains for 5 x 10^6^ generations. Trees were sampled at intervals of 5,000 generations. Two runs were performed and, after confirming convergence of runs, the first 15% of generations were discarded as burn-in, with remaining topologies used to generate a 50% majority-rule consensus. Trees were visualized and edited using iTOL software (Letunic and Bork 2021).

### Pathogenicity test

Jambu seeds were sown in a styrofoam tray of 110 cm³/cell containing Tropstrato HT Hortaliças^®^ substrate (Pine Bark, Vermiculite, PG Mix 14.16.18, Potassium Nitrate, Simple Superphosphate, and Peat – Vida Verde, Mogi Mirim – SP, Brazil) and kept in a growth chamber at 28 °C with daily irrigation. Fourteen days after sowing, the transplant was carried out in plastic pots of 0.5 L, containing a mixture of sand, soil, and substrate in the proportion of 1 : 1 : 1 (v: v: v), previously disinfested using solarization. After 7 days, the seedlings were transplanted into 2 L pots filled with the same mixture used in the 0.5 L pots. Seven days after transplanting, five leaves of each plant were perforated superficially with the aid of a sterilized hypodermic needle and inoculated with *Aphelenchoides* sp. at concentrations of 0 (control), 100, 200, 500, or 1,000 specimens of nematodes/leaf. The inoculation consisted of the deposition of 200 μL of the suspension containing the different concentrations of nematodes on a piece of cotton placed on the adaxial face of previously perforated young leaves of the plant. The volume was completed with the necessary amount of water for the cotton to be completely wet. The control treatment was inoculated with water only. The plants were kept in a greenhouse, with an average temperature of 30 °C ± 5 in a chamber with a sprinkler irrigation system configured to sprinkle water and maintain the relative humidity of the air between 70-80%. After 48 h, the cotton pieces were removed, and the plants were monitored daily for observation and note of symptoms. Leaves were photographed at 2, 6, 15, and 30 days after inoculation (DAI). The experiment was set up in a completely randomized design with 5 treatments and 7 replicates. The experimental unit consisted of a plant with five inoculated leaves.

Thirty DAI, inoculated leaves, and symptomatic leaves (those that were not directly inoculated, but that showed symptoms) were collected, weighed on an analytical balance, and subsequently used for nematode extraction (Coolen and D’Herde, 1972). The final population (Pf) of *Aphelenchoides* sp. was quantified with the aid of a Peters’ chamber under a light microscope. The average number of nematodes in each treatment (P_f_) was used to calculate the reproduction factor (FR), obtained by dividing the P_f_ by the number of inoculated nematodes (P_i_) (Oostenbrink, 1966). In addition, the nematode population was expressed by the number of nematodes g^-1^ in the inoculated leaf and the number of nematodes g^-1^ in the symptomatic leaf.

### Statistical analyzes

The data of FR and Pf were previously submitted to normality (Shapiro –Wilk) and homogeneity of variance of errors (Bartlett) tests. The FR data were transformed by √x to meet these assumptions. Once the assumptions were complied with, the F test was applied through analysis of variance (ANOVA), and the means were compared using Tukey’s test at 0.05 significance. Linear regression analysis was performed for the number of nematodes g^-1^ of the inoculated and symptomatic leaf as a function of the different P_i_ (0, 100, 200, 500, and 1,000 nematodes/leaf).

## Results

### Morphological and morphometric characterization

The morphological and morphometric characters of *Aphelenchoides* sp. are shown in Figure 1 and Table 1, respectively. *Aphelenchoides* sp. had an offset lips, delicate stylet (Fig. 1B), excretory pore generally in front of the posterior portion of the nerve ring (Figs. 1B-C). Presence of four incisures in the lateral field (Fig. 1G). Females, when killed by heat, assume a body slightly arcuate ventrally (Fig. 1A), and males have a “walking-stick” shape. The females had a monodelphic genital tract, and the oocytes present in the growth zone of the ovary were arranged in tandem, oblong spermatheca, usually full of spermatozoa (Fig. 1F). The postvulval sac (PVS) was short (Figs. 1D-E), generally occupying less than a third of the vulva-anus distance (Table 1), but eventually reaching 36.6%, especially in females with PVS containing spermatozoa (Fig. 1E). The tail terminus of the females presented a stellar mucro (with four terminal processes), and in the males, the presence of two terminal processes were generally observed (Fig. 1K). Males had spicules thorn-shaped (Figs. 1H-J; K), with a small condylus, demarcated by a slight depression in the dorsal lamina, while the rostrum was moderately developed with a rounded apex (Figs. 1H-I). Specimens cultured *in vitro*, in general, showed higher mean values for some morphometric characters compared to those derived from plant tissue (Table 1). Despite that, characteristics observed in *Aphelenchoides* sp., in general, overlap with those described for *A. besseyi*, *A. oryzae* and *A. pseudobesseyi* sensu Subbotin et al. (2020), not allowing a safe separation between such species (Table 2).

**Fig. 1.**
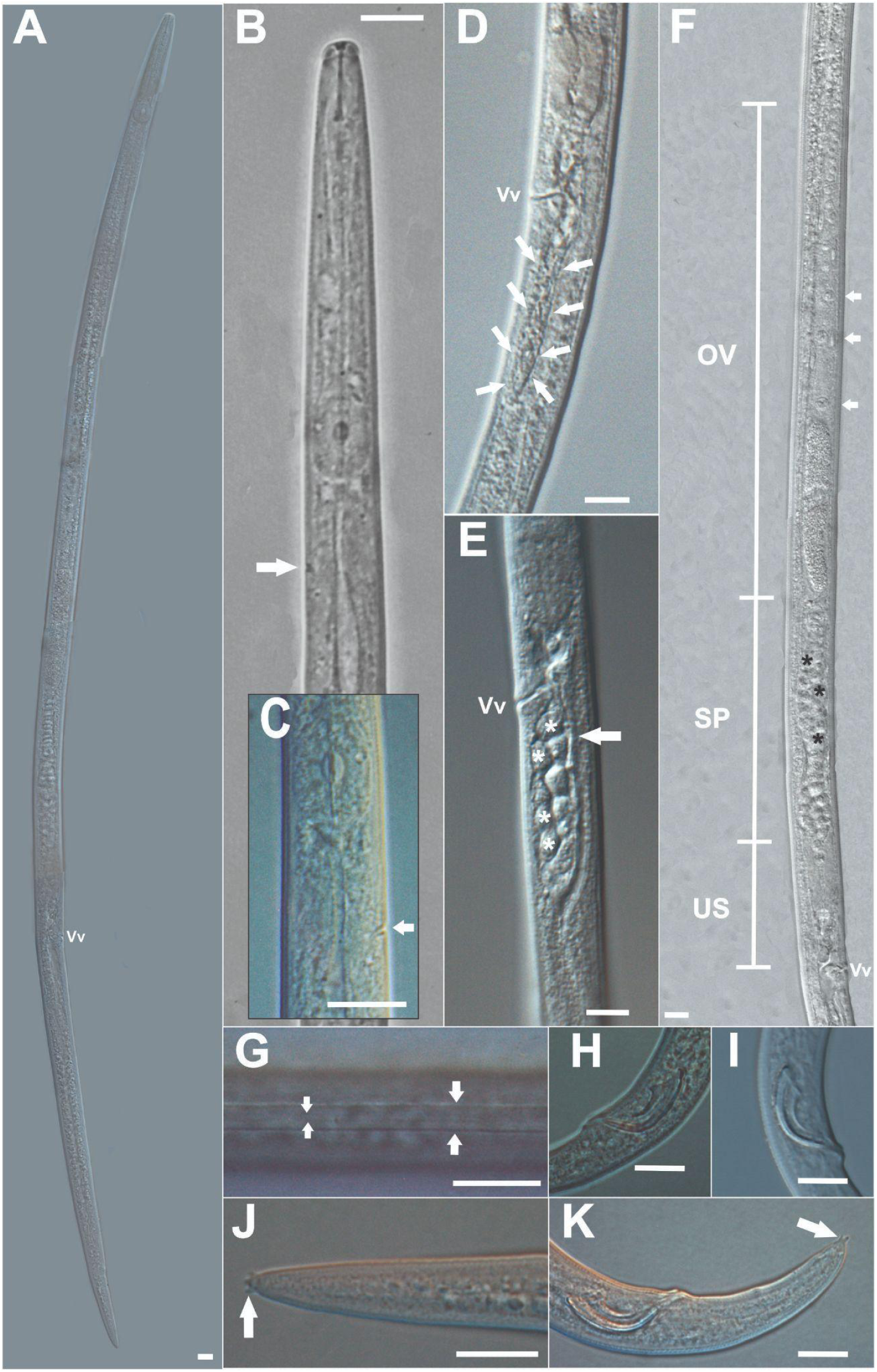
Photomicrographs of females (**A-G; J**) and males (**H-I; K**) of *Aphelenchoides* sp. cultured in *Fusarium pseudocircinatum* **(A-C; F-G; I-K)** or extracted from *Acmella oleracea* leaf naturally infected **(D-E; H). (A):** Entire female body; **(B):** Anterior end region; (**B-C):** The excretory pore (arrowed); Post-vulvar sac (arrowed) **(D)** empty and **(E)** full with spermatozoa (asterisked); **(F):** Female genital tract monodelphic, shown the oocytes in tandem (arrowed) and the spermatheca oblong full with spermatozoa (asterisked); **(G):** Four incisures in the lateral field (arrowed); **(H-I):** Spicules; **(J):** Female tail with tetramucroate caudal end (arrowed); **(K):** Male tail with two terminal processes (arrowed). **Vv**=vulva; **OV**=ovary; **SP**=spermatheca; **US**=uterus. (Bar of scale = 10 µm).

**Table 1.**
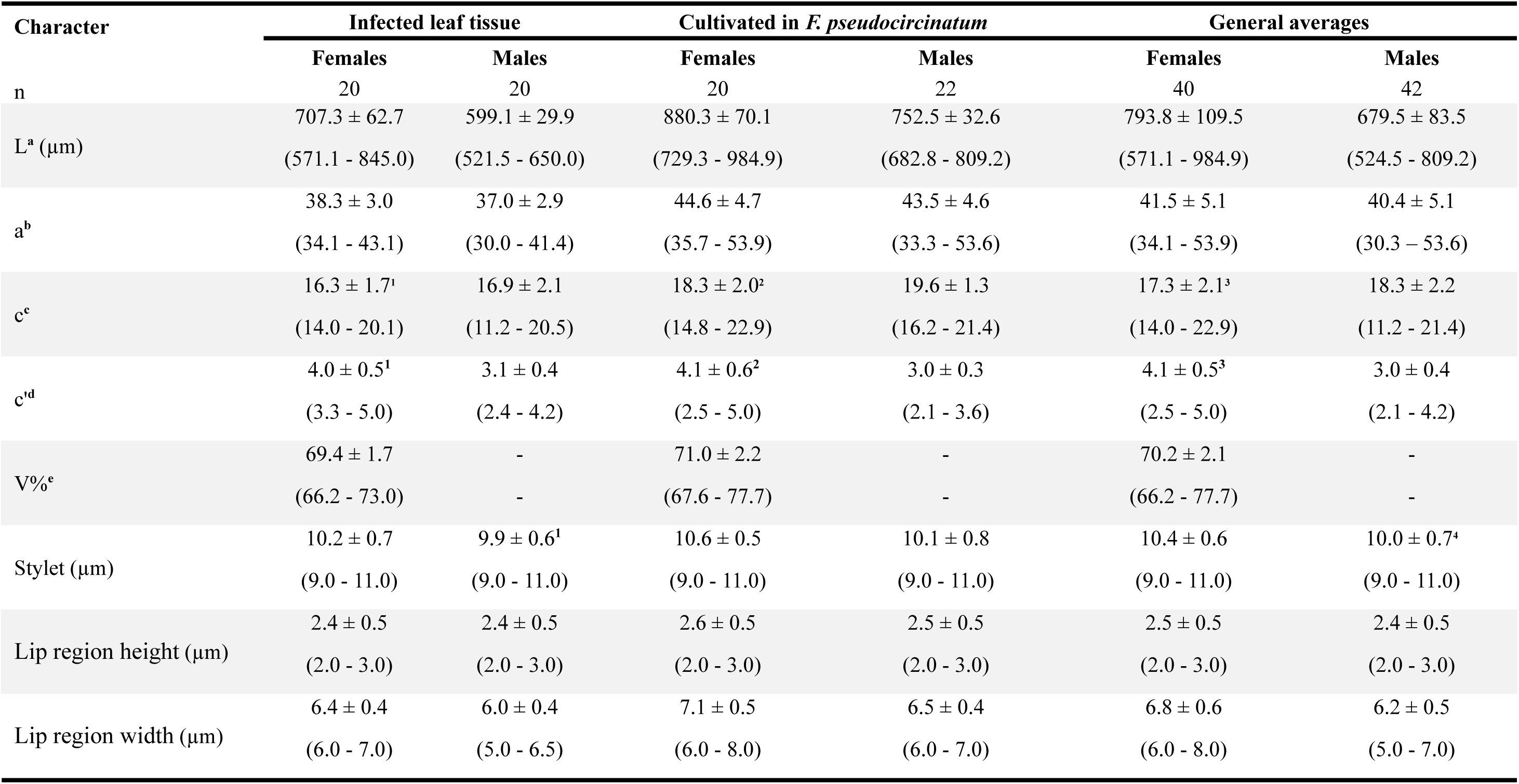
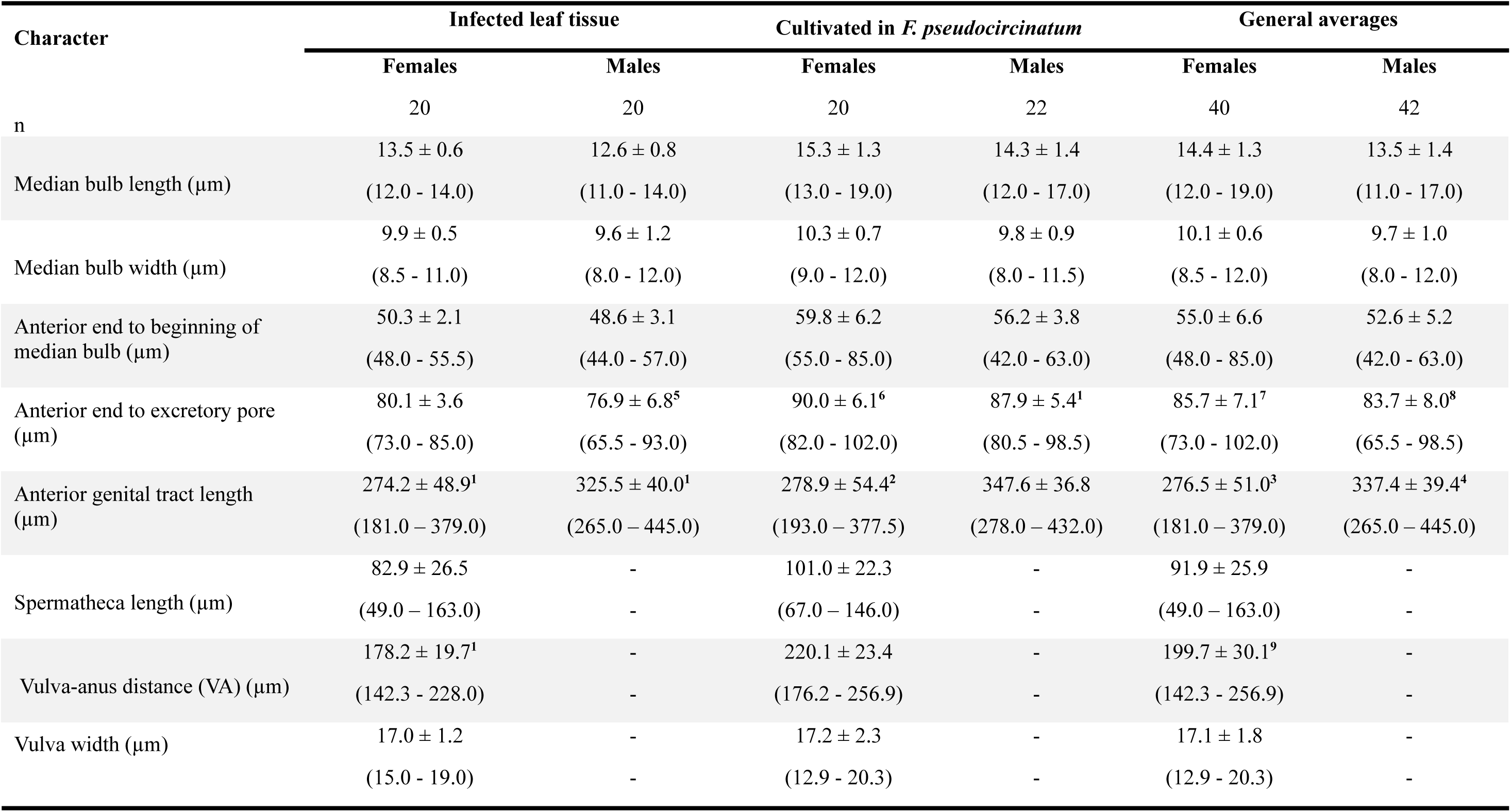
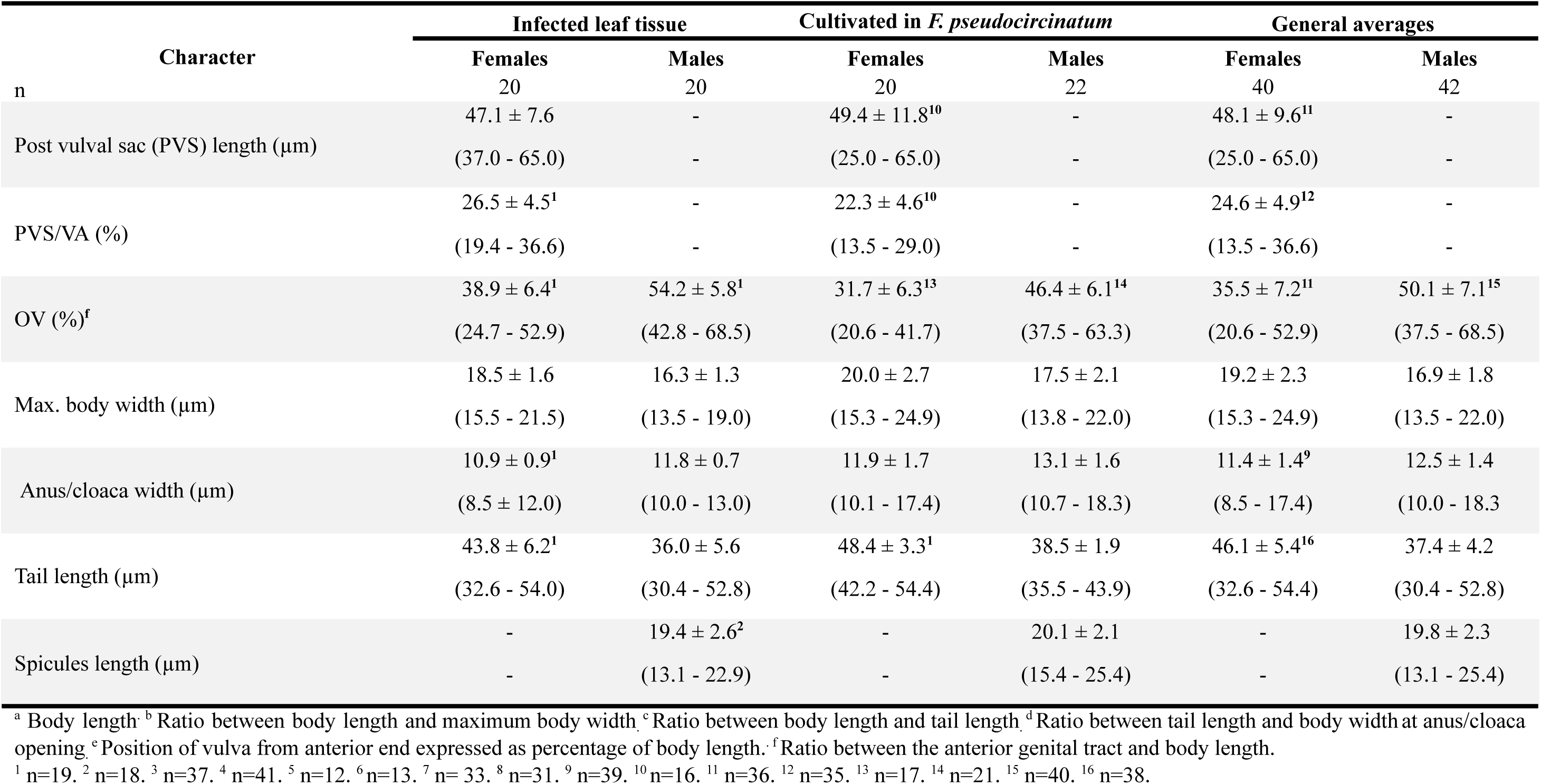
Morphometric characters of *Aphelenchoides* sp. obtained from naturally infected jambu leaf and by *in vitro* multiplication.

**Table 2.** Taxonomic diagnostic characters of *Aphelenchoides* sp. compared to morphologically similar species.

| Species | *L*<br>(μm) | Stylet<br>(μm) | *PVS*<br>/VA% | *a* | *c* | *c'* | Tail<br>(μm) | Tail shape | Excretory pore<br>position | Ovary | Spicules morphology |
| --- | --- | --- | --- | --- | --- | --- | --- | --- | --- | --- | --- |
| *Aphelenchoides*<sup>a</sup><br>sp. (Jambu) | 571-985 | 9-11 | 14-37 | 34-5<br>4 | 14-2<br>3 | 2.5-5 | 33-5<br>4 | Conoid, with 4<br>terminal<br>processes | Usually in front of<br>posterior portion of<br>nerve ring. | Short, oocytes<br>arranged in<br>tandem (single<br>row) | Condylus and rostrum<br>moderately developed |
| *A. besseyi*<br>(Christie, 1942) | 660-750 | n/d<sup>b</sup> | 33 | 32-4<br>2 | 17-2<br>1 | n/d | 36-4<br>2 | Conoid | Slightly anterior to<br>the nerve ring. | Short, oocytes<br>arranged in<br>multiple rows. | n/a |
| *A. besseyi*<br>(Subbotin et al.,<br>2020)<sup>c</sup> | 517-811 | 10.9-1<br>2.5 | 21.9-3<br>3.9 | 34-5<br>3.3 | 14.8-19.1 | 3.6-4.7 | 34-4<br>6 | Conoid, with 3-4<br>terminal<br>processes | Slightly anterior to<br>the nerve ring. | Short, oocytes<br>arranged in<br>multiple rows | Condylus and rostrum<br>clear, not very<br>pronounced |
| *A. pseudobesseyi*<br>(Subbotin et al.,<br>2020)<sup>d</sup> | 540-873 | 11.1-1<br>2.5 | 21.5-4<br>0 | 31.4-49.0 | 14.7-19.6 | 3.4-4.5 | 34-4<br>7 | Conoid, with 2-3<br>terminal<br>processes | Near the anterior<br>end of the nerve<br>ring. | Short, oocytes<br>arranged in 2-4<br>rows | Condylus rounded or<br>rectangular and rostrum<br>moderately developed<br>pointed |
| *A. oryzae* (Yokoo,<br>1948)<sup>e</sup> | 504-732 | 12 | Short | 38-4<br>8 | n/d | n/d | n/d | Conoid, with 3<br>terminal<br>processes | n/a | n/a | n/a |
| *A. oryzae*<br>(Subbotin et al.,<br>2020)<sup>f</sup> | 586-1000 | 11.4-1<br>2.8 | 16.8-2<br>6.1 | 39.7-58.9 | 15.6-22 | 3.6-3.8 | 35-4<br>8 | Conoid, with<br>2-3<br>terminal<br>processes | At the level of the<br>nerve ring | Short, oocytes<br>arranged in 2-4<br>rows | Condylus evident and<br>rostrum pointed |

<sup>a</sup>Minimum and maximum values were obtained from the characters observed in the population extracted from plant tissue and from that multiplied in *F. pseudocircinatum*.
<sup>b</sup>Information not available.
<sup>c</sup>Maximum and minimum values of all populations characterized by Subbotin et al. (2020).
<sup>d</sup>Maximum and minimum values of all populations characterized by Subbotin et al. (2020).
<sup>e</sup>This species described in Japan in 1948 was synonymized with *A. besseyi* by Allen (1952), but its status was revalidated by Subbotin et al. (2020).
<sup>f</sup>Maximum and minimum values of all populations characterized by Subbotin et al. (2020).

### Molecular characterization

Phylogenetic relationship between *Aphelenchoides* sp. and other *Aphelenchoides* species was inferred by analysis of LSU gene sequence datasets. The LSU gene alignment was 827 base pairs (bp) in length. After removing gaps and ambiguous positions, 569 bp was used for phylogenetic analysis. The alignment contained 36 sequences of 9 species of *Aphelenchoides* and *A. fragariae* (KT692710) used as outgroups. Tree topology inferred by Bayesian analysis (Fig. 2) revealed: (i) a highly supported clade containing populations of *A. pseudobesseyi*; (ii) a sister clade, also well supported, containing both sequences of *A. besseyi* and *A. oryzae*; (iii) and other supported 5 clades, containing populations of *A. ritzemabosi*, *A. pseudogoodeyi*, *A. medicagus*, *A. salixae* and *A. fujianensis*, respectively. *Aphelenchoides* sp. sequence from this study (APJ) grouped into the *A. pseudobesseyi* clade, with posterior probability value = 1, along with the 5 sequences (MT271868, MT271869, MT271871 and OQ930285) of *A. pseudobesseyi* and 6 sequences (KT692694, KY510841, KY510840, MH187564, KY510842 and KY510839) previously identified as *A. besseyi* (Fig. 2). The only two sequence types of *A. besseyi* grouped along the *A. oryzae* group, requiring an investigation using additional markers, which might facilitate their separation into two clearly defined and strongly supported branches.

**Fig. 2.**
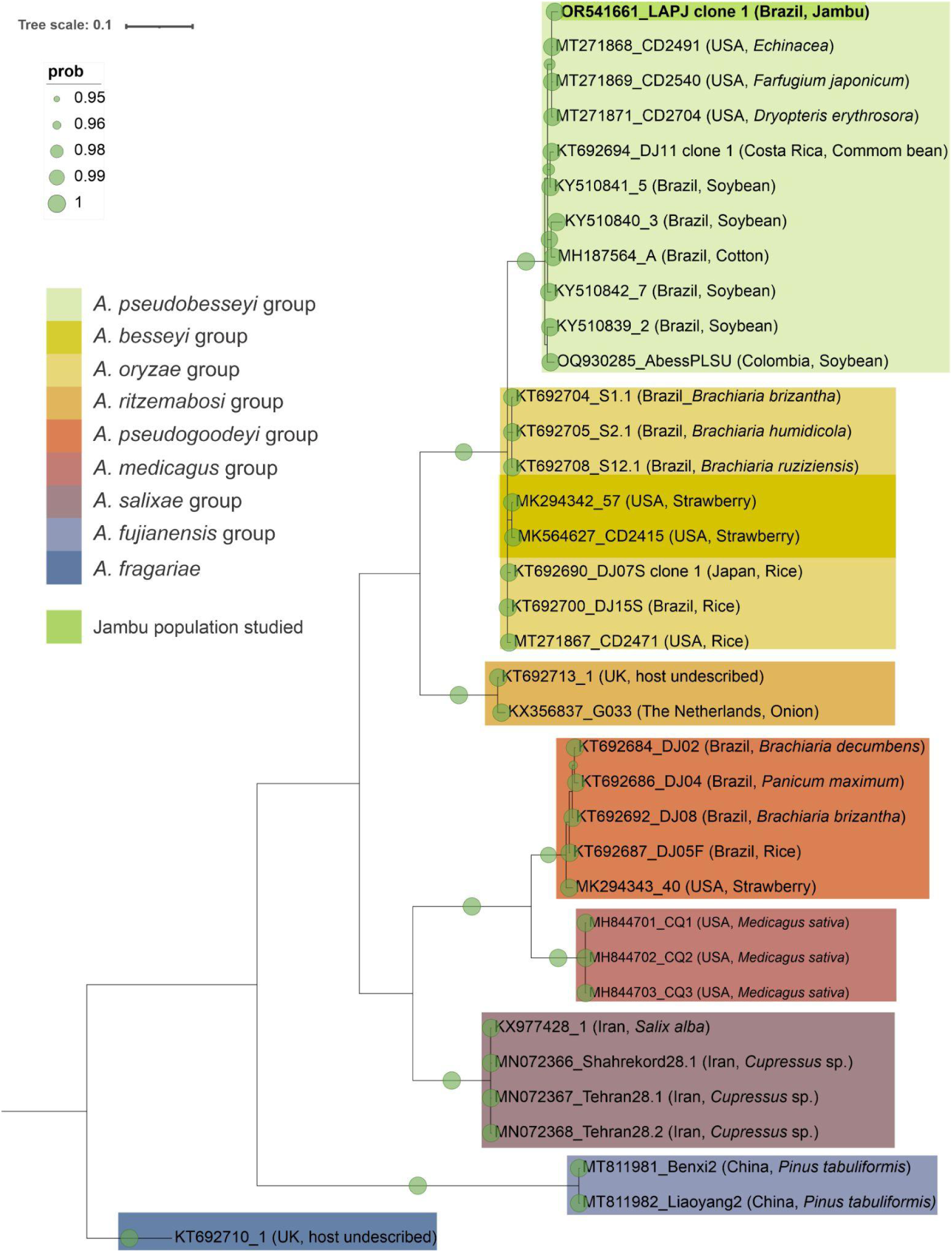
Phylogenetic relationship between species of the genus *Aphelenchoides* as inferred by Bayesian analysis of D2-D3 LSU region of rDNA sequence datasets, with the HKY+I+G model. Posterior probability more than 0.95 are indicated on the branches for respective clades. *A. fragariae* (KT692710) was used as an outgroup species.

### Pathogenicity test

*Aphelenchoides pseudobesseyi* was able to induce symptoms of angular necrotic spots (Figs. 3B-G) and reproduce (Fig. 4; Table 3) in jambu plants of the same genotype originally infected in the field. Symptoms originally described and observed in the field (Fig. 3H) were reproduced under controlled conditions (Fig. 3B-G). When the cotton pieces were removed, after inoculation, angular necrotic lesions were already observed, initially with a soggy appearance due to contact with the cotton humid (Fig. 3B), and with a gradual increase in the severity of the lesions over time, varying according to the initial inoculum concentrations. The angular lesions were initially chlorotic, progressing to necrosis to completely degrade the mesophyll, leaving only the epidermis to maintain the structure of the leaf blade, when under high inoculum pressure (Figs. 3C-D). Such lesions were restricted to the area delimited by the veins, except for the leaves inoculated in the highest inoculum concentration, in which the destruction of a large part of the leaf blade (Fig. 3F). The veins appeared not to suffer damage with the evolution of the symptoms.

**Fig. 3:**
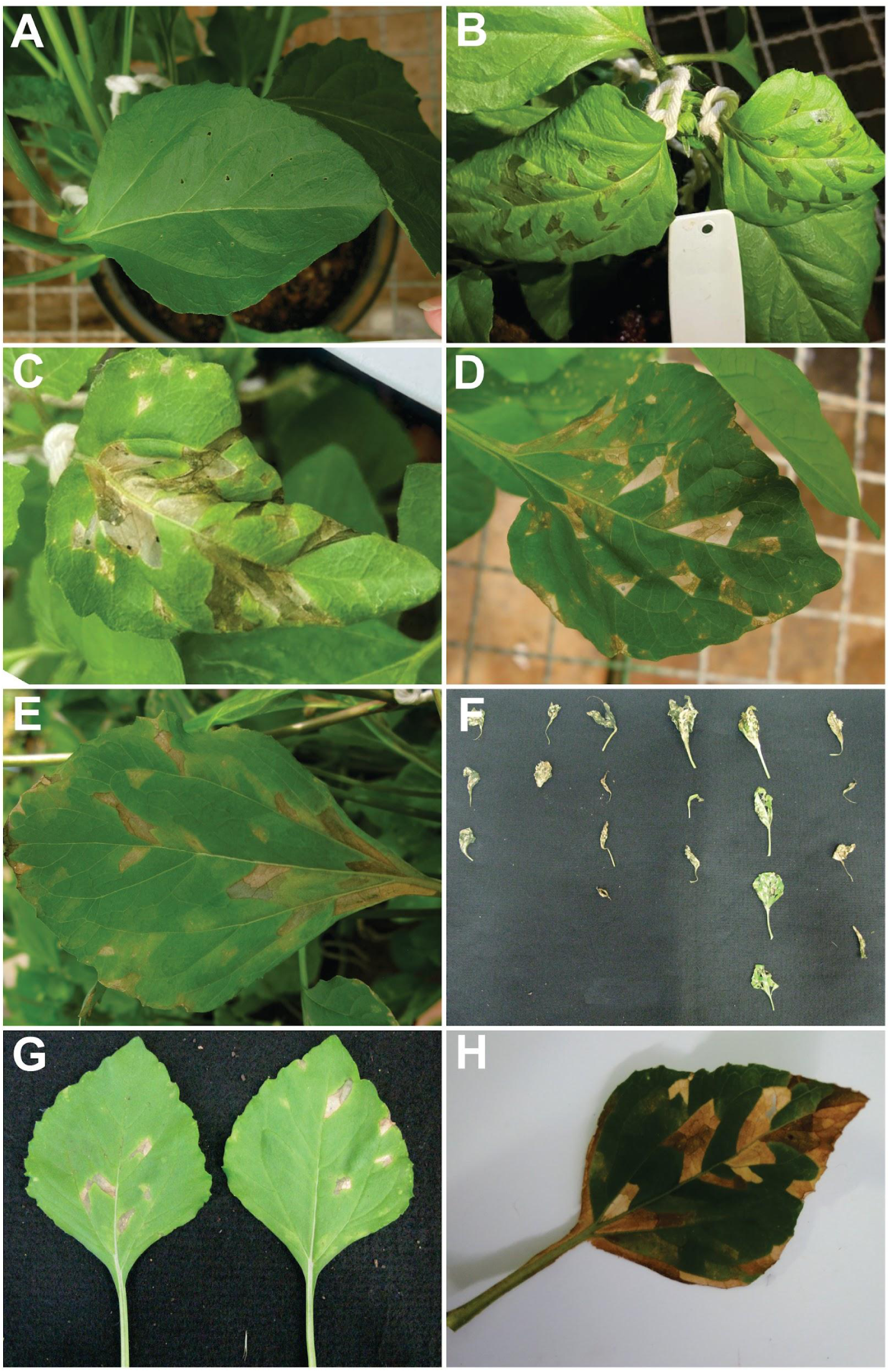
Angular leaf spots induced by *Aphelenchoides pseudobesseyi* in *Acmella oleracea*. **(A)**: Leaf inoculated with water (control) 15 days after inoculation (DAI); **(B)**: Leaves inoculated with 1,000 nematodes, 2 DAI, when initial symptoms started appearing. Note that they were interveinal spots initially soaked and later necrotic; **(C-E):** Necrotic interveinal spots observed at **(C)** 6 DAI, **(D)** 15 DAI and **(E)** 30 DAI inoculated with 1,000, 200 and 1,000 nematode/leaf, respectively; **(F):** Deformation and necrosis of the leaf blade observed at 30 DAI in leaves inoculated with 1,000 nematode/leaf; **(G):** Symptoms of necrotic interveinal spots observed at 30 DAI on leaves adjacent to those inoculated with 1,000 nematodes. This is an evidence that *A. pseudobesseyi* was able to migrate and induce symptoms at sites distant from the site of inoculation; **(H):** Jambu leaf collected in the field, showing the symptoms of necrotic interveinal spots in natural infection caused by *A. pseudobesseyi*.

**Fig. 4:**
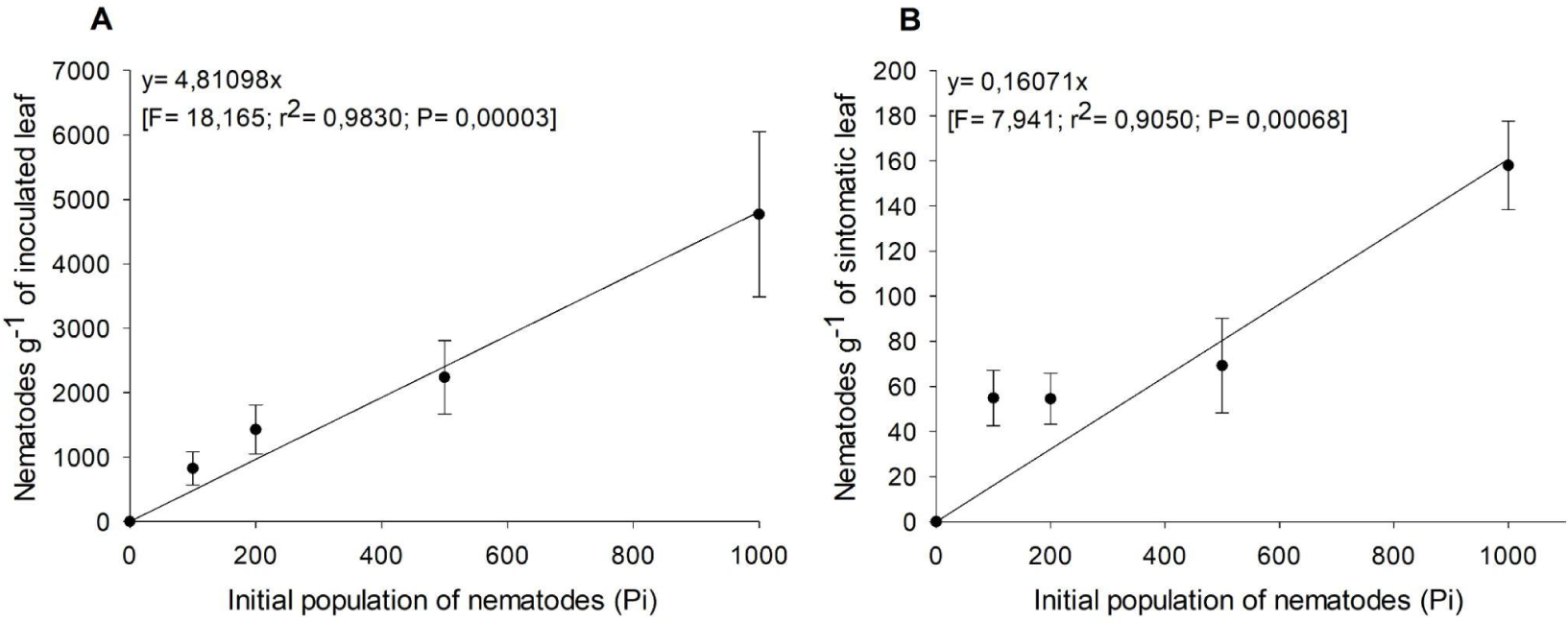
Linear regression of the initial inoculum densities (P_i_) and the final population (P_f_) of *Aphelenchoides pseudobesseyi* **(A):** Extracted directly from inoculated leaves and **(B):** from non-inoculated symptomatic leaves, which were adjacent to those inoculated with increasing inoculum concentrations (100, 200, 500 and 1,000 nematodes/leaf), 30 days after inoculation. Each point represents the mean of seven replicates, and the bars represent the standard deviation.

**Table 3.**
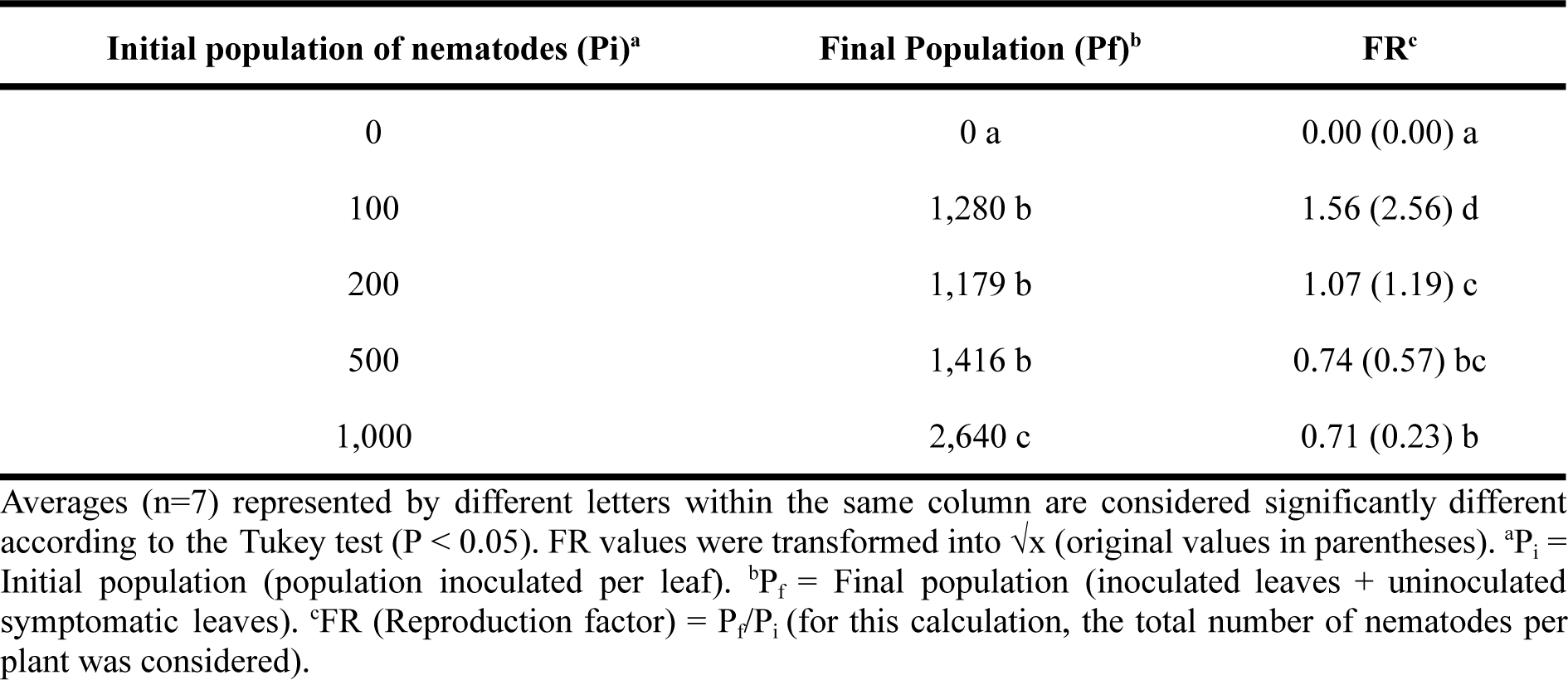
Population density and reproduction factor of *Aphelenchoides pseudobesseyi* obtained 30 days after inoculation.

| Initial population of nematodes ($P_i$)<sup>a</sup> | Final Population ($P_f$)<sup>b</sup> | FR<sup>c</sup> |
| --- | --- | --- |
| 0 | 0 a | 0.00 (0.00) a |
| 100 | 1,280 b | 1.56 (2.56) d |
| 200 | 1,179 b | 1.07 (1.19) c |
| 500 | 1,416 b | 0.74 (0.57) bc |
| 1,000 | 2,640 c | 0.71 (0.23) b |

Averages ( $n=7$ ) represented by different letters within the same column are considered significantly different according to the Tukey test ( $P < 0.05$ ). FR values were transformed into $\sqrt{x}$ (original values in parentheses). <sup>a</sup> $P_i$ = Initial population (population inoculated per leaf). <sup>b</sup> $P_f$ = Final population (inoculated leaves + uninoculated symptomatic leaves). <sup>c</sup>FR (Reproduction factor) = $P_f/P_i$ (for this calculation, the total number of nematodes per plant was considered).

The final population of nematodes obtained at 30 DAI increased proportionally to the initial inoculum density (Fig. 4A). The final population of nematodes obtained with the inoculation of 1,000 nematodes/leaf was almost five times greater than those estimated in the lowest initial inoculum density tested (P_i_ = 100 nematodes/leaf) (Fig. 4A). The highest severity of the symptoms was observed with the inoculation of 1,000 nematodes, resulting in extensive deformation and necrosis of the leaf blade of the inoculated leaves (Fig. 3F). In plants inoculated with nematodes, there was greater leaf abscission than in the control treatment, but there was no clear relationship between this and the amount of inoculated nematodes (data not shown).

Some non-inoculated leaves, adjacent to the ones that were, also showed symptoms from 15 DAI, showing that the nematodes were able to migrate to other leaves and initiate the infective process. In this case, there was an increase in the P_f_ of nematodes in these leaves as the concentration of inoculated nematodes (P_i_) increased (Fig. 4B). There was the reproduction of *A. pseudobesseyi* in jambu leaf tissues. At the two lower initial inoculum concentrations, the FR was higher than 1 (Table 3), being that at P_i_=100 nematodes there was the highest reproduction of *A. pseudobesseyi* in jambu leaves (FR=2.56; P <0.05) (Table 3). In the two higher initial inoculum concentrations (P_i_=500 and P_i_=1,000) the reproduction factors were lower than 1, the outcome of the greater disease severity and tissue destruction (Table 3; Figure 3F).

## Discussion

The genus *Aphelenchoides* harbors hundreds of nematode mycophagous species, with some being facultative plant parasites (Sánchez-Monge et al. 2015). Among the species most harmful to crops, *A. besseyi* Christie, 1942, *A. fragariae* Christie, 1932 and *A. ritzemabosi* Steiner and Buhrer, 1932 deserve to be highlighted. Despite the limitations, morphological and morphometric characterization is still widely used for identifying *Aphelenchoides* species (Chaves et al. 2013; De Jesus et al. 2016; Meyer et al. 2017; Mobasseri et al. 2018). Although useful, the presence of few diagnostic taxonomic characters limits the secure identification of some species of the genus (Jesus and Cares 2016; Oliveira et al. 2019; Subbotin et al. 2020). We identified the causal agent of the jambu angular leaf spot as a species of the *A. besseyi* complex, based on morphological and morphometric characterization.

Recently, the taxonomic status of populations previously identified as *A. besseyi* was reviewed, and evidence supporting the hypothesis of a cryptic species complex was pointed out (Subbotin et al. 2020). Based on morphological analyzes and morphometric measurements combined with phylogenetic and genetic distance analyzes, using different molecular markers, Subbotin et al. (2020) proposed the separation of what until then had been identified as *A. besseyi* into at least three species: *A. besseyi*, *A. oryzae*, and a new species, *A. pseudobesseyi*. The authors highlighted the difficulty in separating these three species only by morphological and morphometric analysis, which all were noted for us.

Thus, the use of molecular markers, mainly those based on three loci of the nuclear ribosomal DNA (LSU, SSU and ITS regions) and mitochondrial DNA (especially the cytochrome oxidase subunit I - COI gene), has been widely employed in taxonomic studies to enhance resolution and reliability (Carta et al. 2019). This combination of molecular and morphological taxonomy is referred to as integrative taxonomy (Jesus et al. 2016; Oliveira et al. 2011; Subbotin et al. 2020).

Regarding these molecular markers, the SSU marker is typically recommended for analyzes at higher taxonomic levels, because of its highly conserved nature. However, it has also demonstrated a robust phylogenetic signal for distinguishing between *Aphelenchoides* species (Rybarczyk-Mydłowska et al. 2012). On the other hand, LSU is recommended for inter-specific analysis because of its balanced blend of conservation and variability (Roberts et al. 2016). It is the most marker used in researches for distinguishing and describing new species within the *Aphelenchoides* genus (Subbotin et al. 2006; Shahabi et al. 2016; Meyer et al. 2017; Buonicontro et al. 2018; Favoreto et al. 2018, Noronha et al. 2020), making it the logical choice for the phylogenetic analysis in this study due to these characteristics.

Despite the robust phylogenetic support observed in the LSU phylogenetic tree generated in this study, among the *A. oryzae* and *A. pseudobesseyi* clades, the type sequences of *A. besseyi* (MK294342 and MK564627) clustered within the *A. oryzae* group, with a low posterior probability value (0.74). As noted in this study and in other research, the morphology of species within the *A. besseyi* complex is highly conserved, with overlapping morphometric characters, which complicates species differentiation, leading them to be characterized as cryptic species (Subbotin et al., 2020). Moreover, the limited number of sequences available for *A. besseyi sensu stricto* or the possibility that the *A. besseyi* complex is still undergoing speciation could justify this grouping between *A. besseyi* and *A. oryzae*. Therefore, the use of other molecular markers such as COI, ITS and SSU, as well as conducting concatenated gene analyzes or even phylogenomic analyzes and species delimitation methods, can enhance our comprehension of the taxonomy within this group. Studies involving populations of the *A. besseyi* species complex from different hosts, evaluating their sexual compatibility and pathogenicity to other hosts, can aid in characterizing this complex of species (Gadagkar et al. 2005; Padial et al. 2010; Hsieh et al. 2012; Janssen et al. 2017; Smythe et al. 2019; Magalhães et al. 2020; Xu et al. 2020).

*Aphelenchoides besseyi sensu lato* causes considerable damage to several crops of economic importance, with emphasis on rice, various ornamental plants, cotton, bean, and soybean (Chaves et al. 2013; Sánchez-Monge et al. 2015; Meyer et al. 2017; Favoreto et al. 2018). As these nematodes have the mycophagous habit, in the absence of plant host species, they can survive in the soil by feeding on fungi present in crop residues. In addition, these nematodes enter anhydrobiosis, which is an ametabolic state that allows survival for long periods under desiccation conditions. These characteristics, associated with the wide range of hosts, make it difficult to manage in areas infested with *Aphelenchoides* spp. (De Jesus and Cares 2016).

*Aphelenchoides besseyi s.l.* has emerged as a phytosanitary problem worldwide, and it is considered among the ten nematodes more damaging to agriculture (Jones et al. 2013). In the United States, this nematode has been reported to cause damage to strawberry plants and ornamental plants, including gerbera (Oliveira et al. 2019). In Costa Rica, *A. besseyi* caused losses from 24 to 70% in common beans crops (Chaves et al. 2013). In Brazil, although there are no reports of *A. besseyi* attacking common beans in commercial areas, most planted cultivars were susceptible to this nematode when inoculated under greenhouse conditions (Favoreto et al. 2018; Favoreto et al. 2021). In addition, *A. besseyi* is reported in the Midwest and Northern regions of Brazil, causing significant losses in soybean (Meyer et al. 2017) and cotton (Favoreto et al., 2018) crops and more recently, identified as inducing leaf spots on yams (*Dioscorea cayenesis* Lam.) in the Northeast region (Noronha et al. 2020). The symptoms observed in soybean and cotton are similar, and included undergrowth, thickening of nodes, flower abortion, leaf deformations, over budding, in addition to delaying the natural senescence of the plants, in the case of soybean (Meyer et al. al. 2017; Favoreto et al. 2018). So far, there are no records of nematodes of the genus *Aphelenchoides* causing damage to the jambu crop.

In this study, we report a new foliar disease in *Acmella oleracea*, the angular spot of the jambu, caused by *A. pseudobesseyi,* occurring in Brazil. To our knowledge, this is the first report of this disease in the world. For future work involving this pathosystem, we recommend inoculation with an initial population of 100 nematodes/leaf.

## Acknowledgements

This study was financed in part by the Coordenação de Aperfeiçoamento de Pessoal de Nível Superior - Brazil (CAPES) - Finance Code 001. The first author received a scholarship from the Fundação de Amparo à Pesquisa do Estado de Minas Gerais - Brazil (FAPEMIG) and from CAPES. The third and fourth authors received scholarships from Conselho Nacional de Desenvolvimento Científico e Tecnológico (CNPq) and CAPES-PROEX, respectively.

## Notes

### Competing Interest Statement

The authors have declared no competing interest.

